# Chronic aromatase inhibition increases ventral hippocampal neurogenesis in middle-aged female mice

**DOI:** 10.1101/434662

**Authors:** Jessica A Chaiton, Sarah J Wong, Liisa AM Galea

## Abstract

Letrozole, a third-generation aromatase inhibitor, prevents the production of estrogens in the final step in conversion from androgens. Due to its efficacy at suppressing estrogens, letrozole has recently taken favor as a first-line adjuvant treatment for hormone-responsive breast cancer in middle-aged women. Though patient response to letrozole has generally been positive, there is conflicting evidence surrounding its effects on the development of depression. It is possible that the potential adverse effects of letrozole on mood are a result of the impact of hormonal fluctuations on neurogenesis in the hippocampus. Thus, to clarify the effects of letrozole on the hippocampus and behavior, we examined how chronic administration affects hippocampal neurogenesis and depressive-like behavior in middle-aged, intact female mice. Mice were given either letrozole (1mg/kg) or vehicle by injection (i.p.) daily for 3 weeks. Depressive-like behavior was assessed during the last 3 days of treatment using the forced swim test, tail suspension test, and sucrose preference test. The production of new neurons was quantified using the immature neuronal marker doublecortin (DCX), and cell proliferation was quantified using the endogenous marker Ki67. We found that letrozole increased DCX and Ki67 expression and maturation in the dentate gyrus, but had no significant effect on depressive-like behavior. Our findings suggest that a reduction in estrogens in middle-aged females increases hippocampal neurogenesis without any adverse impact on depressive-like behavior; as such, this furthers our understanding of how estrogens modulate neurogenesis, and to the rationale for the utilization of letrozole in the clinical management of breast cancer.

## 1. Introduction

Estrogen-suppressive therapy is a common and effective adjuvant treatment of hormone-responsive breast cancer in postmenopausal women. Letrozole, a non-steroidal aromatase inhibitor (AI) that prevents the conversion of androgens into estrogens in the final steps of the estrogen-synthesis pathway, is a first-line treatment of choice. Despite its demonstrated benefits on breast cancer progression, there is conflicting clinical and pre-clinical evidence regarding its adverse effects on mood and cognition. Recently, the effects of letrozole on cognition have attracted more attention, but the evidence for its effects on depression is less understood. Both clinical and pre-clinical trials have found opposing effects of letrozole on mood and behavior (Borbélyová et al., 2017; Chang et al., 2015; Kokras et al., 2018, 2014; Meng et al., 2011). Most animal studies to date have used rodents of varying ages, gonadal hormone status, sex, and duration of treatment, resulting in conflicting data that are poorly understood.

Women are susceptible to developing depression during times of dramatic hormone fluctuations such as postpartum and perimenopause. Suppression of ovarian hormones can induce a depressive-like phenotype in women and rodents (Frokjaer et al., 2015; Mahmoud et al., 2016a), suggesting that a reduction in estrogens renders females more susceptible to depression. Thus, it is possible that the adverse effects of letrozole on mood may be a result of its action on suppressing estrogens.

The hippocampus has a high concentration of estrogen receptors, and is a region that is implicated in the pathoetiology of depression. Estrogens modulate adult hippocampal neurogenesis, with chronic exposure suppressing neurogenesis independent of its effects on upregulating cell proliferation (Mahmoud et al., 2016b). Decreased hippocampal neurogenesis is seen in depressed patients and in animal models of depression, which is restored with antidepressant treatment (Boldrini et al. 2012, Green and Galea, 2008, Mahmoud et al., 2016a). Furthermore, androgens enhance hippocampal neurogenesis in adult male rodents (Hamson et al., 2013) but it is not known whether androgens can modulate neurogenesis in females. Additionally, letrozole modulates cell proliferation in hippocampal dispersion cultures in-vitro from postnatal day 5 rats (Fester et al., 2006). It is possible that changes in neuroplasticity serve as a neural basis for local estrogens to exert their effects on mood. Therefore, we sought to investigate the effects of estrogen suppression due to chronic letrozole treatment on depressive-like behavior and hippocampal neurogenesis in middle-aged female mice.

## 2. Methods

### 2.1 Subjects

Nineteen C57/Bl6J female mice 10-12 months of age were obtained from the Animal Care Centre at the University of British Columbia. All animals were maintained on a 12h light/dark cycle (lights on at 07:00h), group housed (2-3) and given ad libitum access to food (Purina chow) and water. All procedures were performed in accordance with ethical guidelines set by the Canadian Council on Animal Care, and approved by the Animal Care Committee at the University of British Columbia.

### 2.2 Drug preparation and treatment

All animals received daily intraperitoneal (i.p.) injections of 1mg/kg letrozole or saline vehicle for 21 days (see Fig. 1A; dose chosen due to Aydin et al., 2008, Kokras et al., 2014, 2018). Letrozole was dissolved in 0.9% saline at 0.1mg/mL, dissolved with aid of ultrasonic bath. We chose to give i.p. injections rather than oral administration and it is important to acknowledge that route of administration can influence neural and behavioral consequences of drug treatment. However, injections are more likely to give consistent drug quantities compared to oral routes of administration (Kott et al., 2016; Ingberg et al., 2012; Pawluski et al 2014), thereby ensuring accurate dosing.

**Figure 1:**
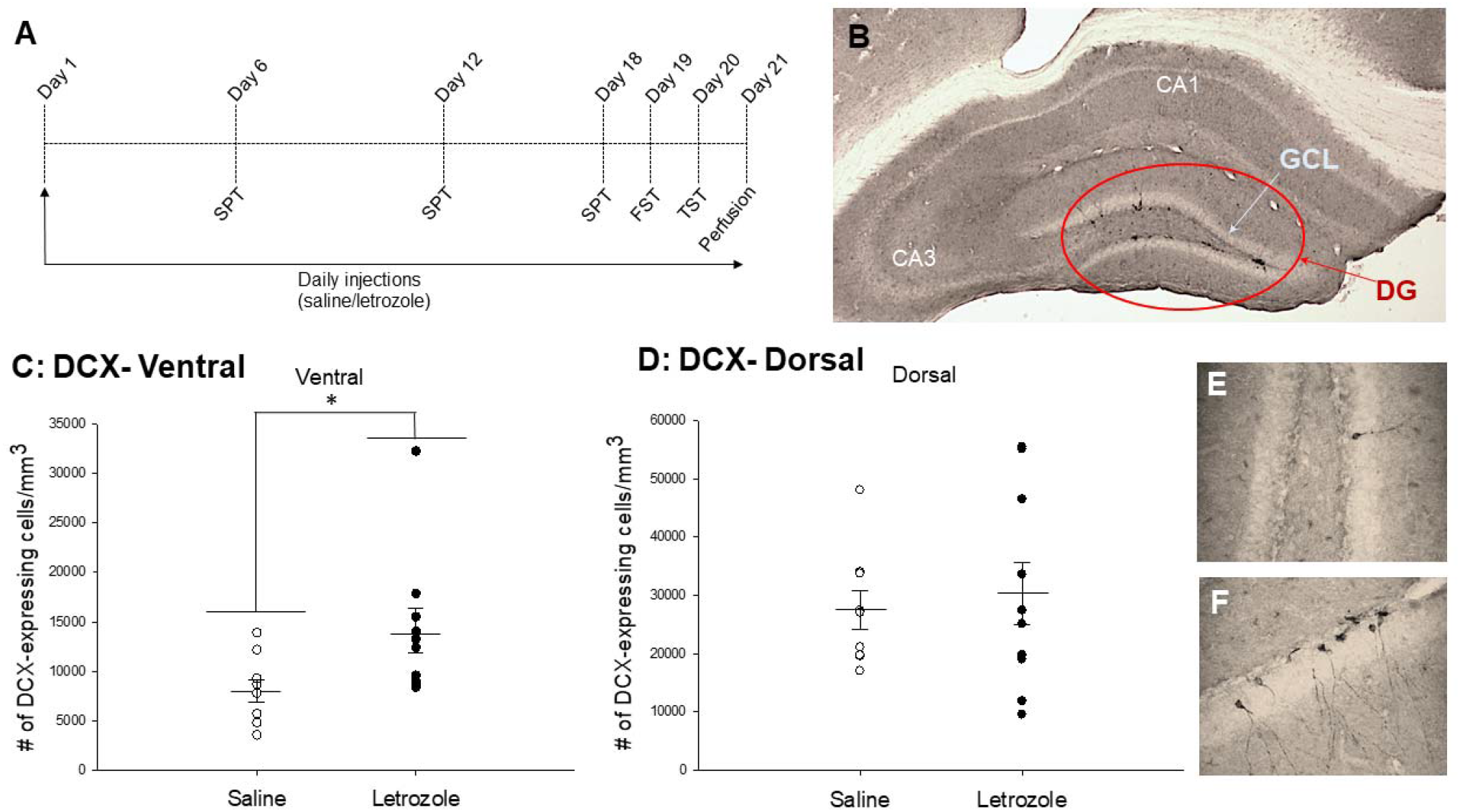
A) Experimental timeline. B) Photomicrograph of the hippocampal region being analyzed viewed at 40x magnification. GCL: granule cell layer. DG: dentate gyrus. C and D) Mean density of doublecortin (DCX)-expressing cells in the dentate gyrus (DG). Letrozole significantly increased the density of DCX-expressing cells in the ventral region (C), but not dorsal (D). E-F) Representative photomicrographs of DCX-expressing cells, viewed at 400X magnification. FST: forced swim test; TST: tail suspension test; SPT sucrose preference test.

### 2.3 Behavioral Testing

Behavioral Testing occurred during days 18-20 of 21 days of letrozole or saline treatment, with the exception of the Sucrose Preference Test, which was administered weekly. A timeline of the procedures is shown in Figure 1A.

#### Forced Swim Test (FST) and Tail Suspension Test (TST)

FST and TST were conducted as described previously (Can et al. 2011; Can et al. 2012; Saeedi Saravi et al., 2016). Each mouse was subjected to a single 6-minute FST and TST session on separate days. FST was conducted in a vertical glass beaker (30cm height x 20cm diameter) filled with clean water (24°C) at a depth of 15cm. In the TST session, mice were suspended by their tails above the ground with a 17cm strip of tape within a 3-walled rectangular chamber. Both tests were videotaped and scored using BEST collection software (Educational Consulting, Hobe Sound, FL, USA) by an individual blind to treatment condition. Percent time spent in mobile and immobile behaviors were analyzed, excluding the first two minutes (Can et al., 2011).

#### Sucrose Preference Test (SPT)

Each mouse was habituated to a 1% sucrose solution and the two-bottle procedure by introducing two identical bottles with water and 1% sucrose (counterbalanced) into their home cage for a 48h period. After acclimatization, the test was administered for three days before the start of treatment (baseline) and then once a week for 3 weeks over the course of letrozole treatment as previously done (Gross and Pinhasov, 2016; Mahmoud et al. 2016; Strekalova et al. 2006; Wainwright et al. 2016). Briefly, mice were single housed and simultaneously food and water deprived for 4h. Mice were then presented with 2 bottles for 12h between 20:00h and 08:00h, after which they were re-paired with cage mates. Sucrose preference was calculated using the formula: sucrose preference=(sucrose consumed/(sucrose + water consumed))x100.

### 2.4 Tissue Collection

Twenty-four hours after TST, mice were given an overdose of sodium pentobarbital, and blood was collected by cardiac puncture. Mice were transcardially perfused with 0.9% saline followed by 4% paraformaldehyde. Brains were extracted and post-fixed in paraformaldehyde overnight at 4°C. Brains were transferred to 30% sucrose and stored at 4°C. Brains were sliced in 30μm coronal sections using a Leica SM2000R microtome (Richmond Hill, Ontario, Canada). Sections were stored in antifreeze (20% glycerol and 30% ethylene glycol in 0.1M PBS) at – 20°C until processing.

### 2.5 Doublecortin (DCX) Immunohistochemistry

Sections were rinsed in phosphate buffered saline (PBS) and treated with 0.6% hydrogen peroxide in dH20 for 30 minutes. Sections were rinsed and incubated for 24h at 4°C in primary antibody solution: 1:1000 goat anti-doublecortin (Santa Cruz Biotechnology, Santa Cruz, CA, USA), 0.04% Triton-X in PBS, and 3% normal rabbit serum. Sections were then rinsed and incubated in secondary antibody solution for 24h at 4°C: 1:1000 rabbit anti-goat (Vector Laboratories, Burlington, ON, Canada) in 0.1M PBS. Then, sections were rinsed and incubated in an avidin-biotin complex (ABC Elite Kit, 1:1000, Vector Laboratories) in PBS for 2hr. Sections were rinsed and subsequently 2 x 2min in 0.175M sodium acetate buffer. Immunoreactants were visualized using diaminobenzadine (DAB) in the presence of nickel (DAB peroxidase substrate kit, Vector), mounted on slides, dried, dehydrated and coverslipped.

### 2.6. Ki67 Immunohistochemistry

Sections were rinsed in phosphate buffered saline (PBS) and treated with 0.6% hydrogen peroxide in dH2O for 30 minutes. Sections were rinsed and incubated for 45 minutes at 90°C in 2X saline-sodium citrate buffer. Sections were rinsed with PBS and then incubated for 1 hour at room temperature in 2% normal horse serum and 0.2% Triton-X in PBS. Sections were rinsed with PBS and incubated for 20 hours at 4°C in primary antibody solution 1:3000 mouse anti-Ki67 (BD Biosciences, San Jose, CA, USA), 2% normal horse serum, and 0.2% Triton-X in PBS. Sections were then rinsed with PBS and incubated for 1 hour at room temperature in secondary antibody solution 1:1000 anti-mouse (Vector Laboratories, Burlington, ON, Canada), 2% normal horse serum, and 0.1% bovine serum albumin in PBS. Sections were then rinsed in PBS and incubated for 1 hour at room temperature in avidin-biotin complex (ABC Elite Kit, 1:500, Vector Laboratories, Burlington, ON, Canada) in PBS. Sections were rinsed with PBS. Immunoreactants were visualized using diaminobenzidine (DAB peroxidase substrate kit, Vector) and rinsed with PBS. Sections were then mounted on slides, dried, dehydrated, and coverslipped.

### 2.7 Microscopy, cell quantification, and cell phenotyping

An investigator blinded to treatment condition quantified DCX- and Ki67-expressing cells. DCX-expressing cells were quantified in the granule cell layer of the dentate gyrus in every 10^th^ section along the rostral-caudal axis, using the 40x objective on an Olympus CX22LED brightfield microscope. Raw counts were multiplied by 10 to get an estimate of the total number of DCX-expressing cells, separately in dorsal and ventral regions. Ki67-expressing cells were quantified in the granule cell layer of two dorsal and two ventral slices along the rostral-caudal axis, using the 100x objective on a Nikon E600 microscope. Areas of the granule cell layer of each slice counted were quantified using ImageJ (NIH, Bethseda, MD) and used for density calculations (number of cells per mm^3^).

DCX morphology (Figure 2A) was analyzed using the 100× objective on an Olympus CX22LED brightfield microscope. 50 DCX-expressing cells (25 dorsal GCL and 25 ventral GCL) were randomly selected for each animal, and categorized into one of three maturational stages based on previously established criteria (Plumpe et al. 2006): proliferative (no process or short process), intermediate (medium process with no branching), or post-mitotic (long processes with branching into the GCL and molecular layer).

**Figure 2.**
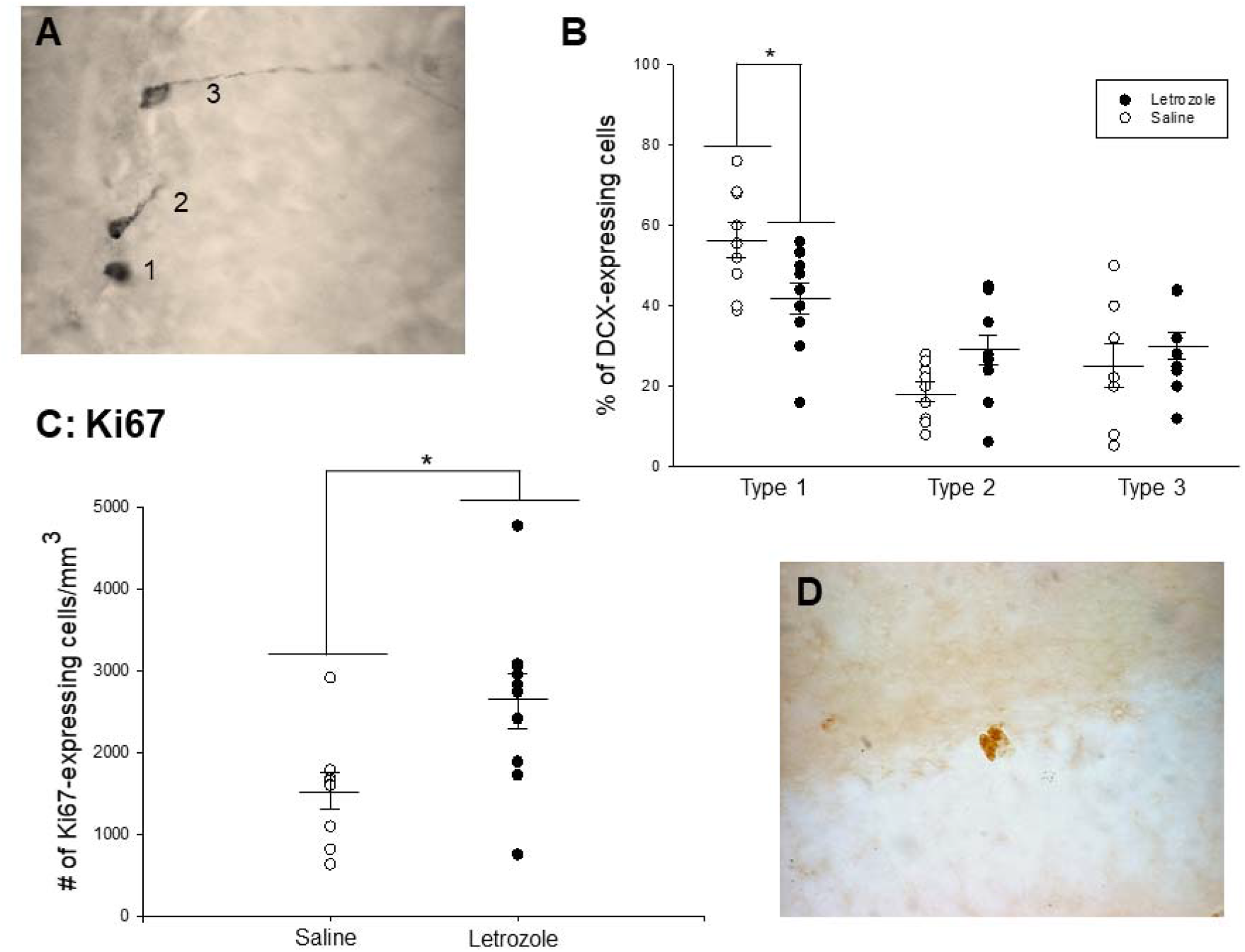
A) Photomicrograph at 1000x of the three types of DCX-expressing cells based on morphology: Type 1 cell, proliferative; Type 2 cell, intermediate;Type 3 cell, post-mitotic, modified definitions from Plümpe et al., 2006. B) Percentage of DCX-expressing cells in the DG in each maturational stage. Letrozole decreased the proportion of type 1 (proliferative) DCX-expressing cells compared to controls, with a non-significant increase the proportion of type 2 (intermediate) DCX-expressing cells (p<0.06). C) Mean density of ki67-expressing cells in the dentate gyrus. Letrozole significantly increased the total density of ki67-expressing cells. D) Photomicrograph taken at 1000x of a cluster of Ki67-expressing cells. *p<0.05. ± SEM.

### 2.8 Determination of estrous cycle phase and serum 17β-estradiol levels

Vaginal cells were collected by lavage on days 17-21 of the experimental timeline. Estrous cycle phase was determined as described previously (Brummelte and Galea, 2010).

Serum 17β-estradiol was quantified by radioimmunoassay kit according to manufacturer’s instructions (DSL-4800 Ultra-sensitive Estradiol RIA, Beckmann Coulter, Missisauga, ON).

### 2.9 Data Analyses

All statistical analyses were performed using Statistica software (Tulsa, OK). Behavioral tests (TST, FST), density of Ki67- and DCX-expressing cells, and morphology of DCX-expressing cells were each analyzed using repeated measures analysis of variance (ANOVA) with drug treatment (letrozole or vehicle) as the between-subjects factor and behavior (immobile, mobile), region (dorsal, ventral) or cell type (type 1,2,3) as within-subjects factor with age as a covariate. Serum 17β-estradiol levels, uterine mass and adrenal mass were analyzed using a student’s t-test. Percent sucrose preference and percent change in body mass were analyzed with a repeated measures ANOVA with drug treatment as the between-subjects factor and week as the within-subjects factor. Post-hoc analyses used the Newman-Keuls test, and a priori tests utilized Bonferroni corrections. We also tested for violations of normality (Kolmogorov-Smirnov test) and homogeneity of variance for each variable. These assumptions were not violated, so parametric statistics were conducted. Pearson product moment correlations were also conducted on variables of interest.

## 3. Results

### 3.1 Letrozole upregulated the density of immature neurons (DCX-expressing cells) and cell proliferation (Ki67-expressing cells) in the dentate gyrus

Letrozole treatment significantly increased the density of DCX-expressing cells (main effect of treatment (F(1,16)=8.12, p<0.011, 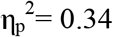) in the granule cell layer of the dentate gyrus (Figure 1C). Indeed, letrozole treatment increased the density of DCX-expressing cells in the ventral (p=0.004; cohen’s d=1.09; Figure 1C) more so than the dorsal region (p=0.06, Cohen’s d=0.20; Figure 1D). Overall there were more DCX-expressing cells in the dorsal compared to the ventral region (main effect of region: p=0.02, 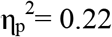).

Letrozole treatment increased the number of Ki67-expressing cells, regardless of region (main effect of treatment F(1,17)=6.28, p=0.023, 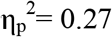; Figure 2C). There were no other significant main or interaction effects (p’s>0.3). Because proestrous state can increase cell proliferation (Tanapat et al., 1999) we ran a t-test to determine if proestrous state was associated with increased Ki67-expressing cells but this was not significant (t(17)=1.59, p=0.13).

### 3.2 Letrozole decreased the proportion of proliferative DCX-expressing cells and increased the proportion of more mature DCX-expressing cells in the ventral dentate gyrus

A priori analysis revealed letrozole decreased the percentage of proliferative (type 1, Figure 2B) DCX-expressing cells compared to saline in the ventral region (p<0.012; Cohen’s d= 1.17) but not the dorsal region (p=0.77; Cohen’s d=0.12). There was a non-statistically significant increase in the proportion of intermediate (type 2, Figure 2B) DCX-expressing cells (p=0.06) in the letrozole-treated mice, but no other statistically significant differences between groups (p’s >0.2).

### 3.3 Letrozole had no significant effect on behavior in the SPT, TST, and FST

In the FST, TST, or SPT there were no significant differences in behaviors between groups (all p’s > 0.15, Figure 3A-C).

**Figure 3.**
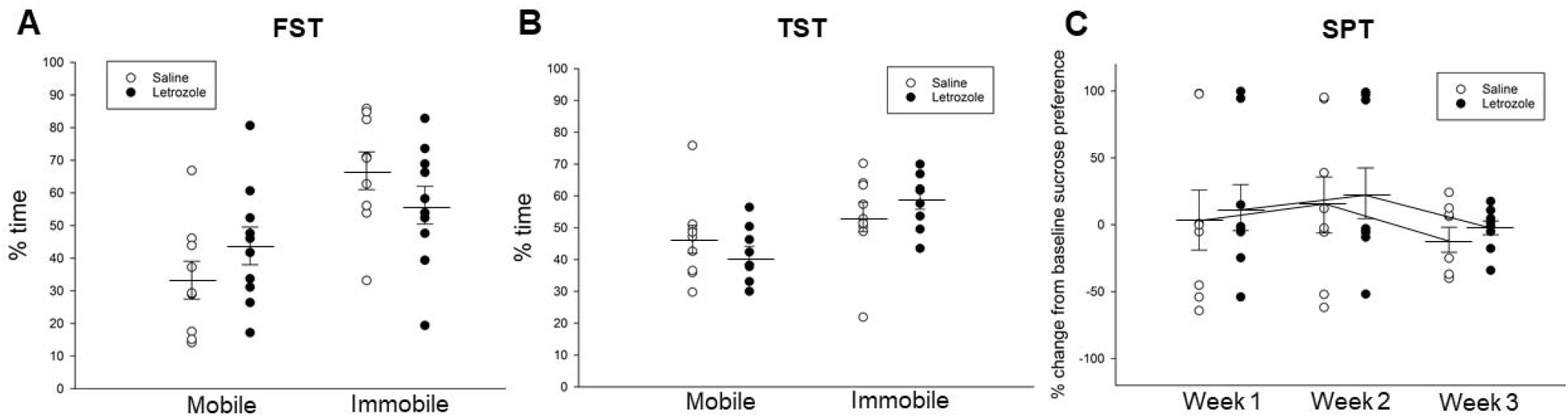
**A-C:** Letrozole had no significant effect on any measure of depressive-like behavior (all p’s > 0.15). A) Percent time spent mobile and immobile in the forced swim test. B) Percent time spent mobile and immobile in the tail suspension test. C) Percent change from baseline sucrose preference over 3 weeks of the sucrose preference test. ±SEM.

### 3.4 Letrozole significantly decreased uterine mass and serum 17β-estradiol levels, but did not influence estrous cycle

Letrozole-treated mice had significantly lower uterine mass than controls (t(17)=2.55, p=0.02; Cohen’s d=0.966; Table 1). Consistent with this outcome, letrozole decreased 17 β-estradiol levels (letrozole: 14.4 ± 1.5) compared to controls (18.9±1.9; t(8)=1.78, p=.05, one-tailed, Cohen’s d=1.15). The values of estradiol were low likely due to age. There was no significant effect of letrozole on body mass from baseline to the last day or adrenal mass (p’s > 0.41; Table 1). In the saline group, 6 mice were in constant diestrus, 2 in estrus and 1 was cycling, while in the letrozole group 4 were in constant diestrus, 4 in constant estrus and 2 were cycling. These distributions were not significantly different according to a chi-square (χ^2^=1.35, p=0.51). There were no significant effects of letrozole to influence body or adrenal mass (p’s < 0.11; Table 1).

**Table 1.** Serum 17β-estradiol concentrations, body, and organ mass across both groups. Letrozole decreased uterine mass and serum 17β-estradiol levels.

|  | Saline | Letrozole |
| --- | --- | --- |
| **Uterine mass (g)** | $0.06 \pm 0.005$ | $0.045 \pm 0.005^*$ |
| **Body Mass (g) baseline** | $26.3 \pm 0.66$ | $25.65 \pm 0.81$ |
| **Body Mass (g) last day** | $26.3 \pm 0.55$ | $26.05 \pm 0.95$ |
| **Adrenal mass (g)** | 0.0063±0.0008 | 0.007±0.0002 |
| **17β-estradiol levels (pg/ml)** | 18.9±1.9 | 14.4± 1.5* |

\* Significantly different from saline-treated groups.

### 3.5 Correlations

There were no significant correlations between neurogenesis markers and behavior (P’s >0.3).

## 4. Discussion

We found that letrozole increased the density of immature neurons in the ventral dentate gyrus of middle aged females. This is consistent with findings that long-term ovariectomy in middle-aged female rats increased, while estrogens decreased the survival of immature neurons (Barha et al., 2015). Additionally, we found that letrozole lowered the proportion of the least mature DCX-expressing neurons. This suggests that letrozole increased the rate of maturation of immature neurons, promoting survival past the proliferative phase into the intermediate and postmitotic phase, consistent with a trend for letrozole to increase type 2 DCX-expressing cells. As DCX is expressed for 28 days in mice (Snyder et al., 2009), this would have entailed that some of the more mature cells were produced without letrozole exposure. However, when we examined cell proliferation using the endogenous marker Ki67, we found that letrozole increased cell proliferation in the dentate gyrus, regardless of region. Ki67 is expressed for approximately 24 h in every part of the cell mitotic cycle except for G_0_ and the initial stages of G_1_. The discrepancy between the length of expression of Ki67 and DCX (1 day and up to 28 days, respectively), likely accounts for the opposing effects seen on DCX-expressing type 1 cells versus Ki67-expressing, as type 1 cells likely expressed DCX for a longer timeframe. It is also important to acknowledge that Ki67 may have been influenced by TST, whereas type 1 DCX cells could have been influenced throughout the last 4 days of behavior testing. It is also possible that the stress effects associated with behavior testing could influence Ki67 expression. However, females do not normally show a decrease in cell proliferation in response to acute stress (Falconer and Galea, 2003, Tzeng et al., 2014), unlike males (Falconer and Galea, 2003, Tzeng et al., 2014). Finally, we find no significant correlations between Ki67-expression and immobility in the TST or FST. The lack of association between variables suggests that the stress of testing did not influence Ki67 expression in middle-aged female mice.

Studies from the Rune laboratory have found that letrozole decreases cell proliferation in hippocampal cultures taken from postnatal day 5 Wistar rats (Fester et al., 2006) and decreases synapses in immature (postnatal day 9) and adult intact female Wister rats (Bender et al., 2010). These findings are consistent with our findings that letrozole decreased the percentage of proliferative (Type 1) DCX-expressing cells, but inconsistent with our results that chronic letrozole increased cell proliferation (Ki67-expressing cells). However, these findings taken all together, suggest that there are age and possibly sex-related differences in the effects of letrozole on cell proliferation and neurogenesis in the hippocampus. Future studies should determine the impact of chronic letrozole on the survival of new neurons independent of its effects on cell proliferation, in an age- and sex-dependent manner.

Interestingly, letrozole’s effects on the density of DCX-expressing cells were seen exclusively in the ventral hippocampus. The ventral hippocampus is implicated in modulating stress and affect (Fanselow and Dong, 2010). Curiously, survival of new neurons in females is increased in the ventral hippocampus with trace eyeblink conditioning involving a shock (Dalla et al. 2009), but not after pattern separation (Yagi et al., 2016) or Morris water maze training (Chow et al., 2013). It would be interesting to examine whether the increase in cell proliferation with letrozole affects fear or appetitive-based conditioning and cognition. Given this, coupled with our lack of findings on affective behavioral measures, it suggests neural consequences of letrozole may be seen either prior to any affective behavioral changes, or on different types of behavior with more prolonged letrozole exposure.

Chronic letrozole had no significant effect on measures of depressive-like behavior (FST, TST, SPT) in the present study, consistent with other studies examining immobility in the FST (Kokras et al., 2018, 2014; Meng et al., 2011). However, acute letrozole in young, ovariectomized female rats had an antidepressant effect in the FST, while chronic letrozole had no such effect (Kokras et al., 2014). Chronic letrozole in young, ovariectomized mice increased anxiety in the open field test and elevated plus maze (Meng et al., 2011) while chronic letrozole in middle-aged cycling female rats had no effect on anxiety in the elevated plus maze (Borbélyová et al., 2017). Collectively these results suggest that chronic letrozole treatment has no significant effect on depressive-like behavior in middle-aged females. Our values in immobility in the FST were high even for saline-injected females (~60%) which is much different from control values for FST in female rats of approximately 20% immobility (Mahmoud et al., 2016). However, our findings are consistent with data in mice with studies indicating levels of immobility in untreated mice around 50-90% of immobility (Akanmuet al 2007; Can et al., 2012; Frye, 2011, Liou et al., 2012). There are known differences between mice and rats in the FST (Molendijk and de Kloet ER, 2019) but to our knowledge no one has described the differences in baseline immobility time as a function of mice versus rats. In the future, the issue of interpretation of the FST would benefit from a careful review of the literature on FST findings in mice versus rats. Future studies should consider the effects of letrozole on depressive-like behavior in the context of a challenge such as chronic stress or cancer, because situations that increase inflammation could impact the effects of letrozole and may provide more robust effects on behavior.

It is possible that an increase in testosterone that accompanies aromatase inhibition is responsible for mitigating negative behavioral effects of estrogen depletion. We did attempt to measure serum testosterone in this study, but the levels in serum were below detection threshold in middle-aged females and future studies should examine testosterone levels in brain after chronic letrozole treatment in middle-aged female mice. Castrated male rats are more susceptible to developing depressive-like endophenotypes following chronic stress (Wainwright et al., 2011). Similarly, in men, hypogonadism is associated with depressive symptoms, and androgen treatment can ameliorate these symptoms (Zarrouf et al., 2009). Furthermore, testosterone given to surgically menopausal women can elevate mood (Sherwin, 1988). Further studies could evaluate the role of testosterone on mood after aromatase inhibition in middle-aged females. Finally, in our study we saw no significant effects of letrozole on depressive-like or coping strategies in middle-aged females, but other studies have seen effects of acute letrozole on memory in female mice (Tuscher et al., 2016) and the testing of the effects of chronic letrozole on decision making may be warranted.

## 5. Conclusion

We demonstrate that in middle-aged female mice, chronic letrozole increased hippocampal neurogenesis but had no effect on depressive-like behavior. Future studies could investigate more behaviors that may be regulated by ventral neurogenesis in the hippocampus such as fear motivated learning (Dalla et al., 2009). Furthermore, it is important for studies to investigate the mechanisms behind potential neuropsychiatric effects of aromatase inhibitors on middle-aged women, which may shed light on the impact of adjuvant cancer treatments on quality of life.

## Acknowledgements

We gratefully acknowledge funding from Canadian Institutes of Health Research (MOP 142308) to LAMG and a Faculty of Medicine award (Summer Student Research Program) at University of British Columbia for salary to JC.

## References

Aydin, M., Yilmaz, B., Alcin, E., Nedzvetsky, V.S., Sahin, Z., Tuzcu, M., 2008. Effects of letrozole on hippocampal and cortical catecholaminergic neurotransmitter levels, neural cell adhesion molecule expression and spatial learning and memory in female rats. Neuroscience 151, 186–194. https://doi.org/10.1016/j.neuroscience.2007.09.005

Barha, C.K., Lieblich, S.E., Chow, C., Galea, L.A.M., 2015. Multiparity-induced enhancement of hippocampal neurogenesis and spatial memory depends on ovarian hormone status in middle age. Neurobiol. Aging 36, 2391–2405. https://doi.org/10.1016/j.neurobiolaging.2015.04.007

Bender RA1, Zhou L, Wilkars W, Fester L, Lanowski JS, Paysen D, König A, Rune GM. Roles of 17ß-estradiol involve regulation of reelin expression and synaptogenesis in the dentate gyrus. Cereb Cortex. 2010 Dec;20(12):2985–95.

Boldrini, M., Hen, R., Underwood, M.D., Rosoklija, G.B., Dwork, A.J., Mann, J.J., Arango, V., 2012. Hippocampal Angiogenesis and Progenitor Cell Proliferation Are Increased with Antidepressant Use in Major Depression. https://doi.org/10.1016/j.biopsych.2012.04.024

Borbélyová, V., Domonkos, E., Csongová, M., Kačmárová, M., Ostatníková, D., Celec, P., Hodosy, J., 2017. Sex-dependent effects of letrozole on anxiety in middle-aged rats. Clin. Exp. Pharmacol. Physiol. 44, 93–98. https://doi.org/10.1111/1440-1681.12731

Brummelte, S., Galea, L.A.M., 2010. Chronic high corticosterone reduces neurogenesis in the dentate gyrus of adult male and female rats. Neuroscience 168, 680–690. https://doi.org/10.1016/J.NEUROSCIENCE.2010.04.023

Can, A., Dao, D.T., Terrillion, C.E., Piantadosi, S.C., Bhat, S., Gould, T.D., 2012. The tail suspension test. J. Vis. Exp. e3769. https://doi.org/10.3791/3769

Can, A., Dao, D.T., Arad, M., Terrillion, C.E., Piantadosi, S.C., Gould, T.D., 2011. The Mouse Forced Swim Test. J. Vis. Exp. e3638. https://doi.org/10.3791/3638

Chang, C.-H., Chen, S.-J., Liu, C.-Y., 2015. Adjuvant treatments of breast cancer increase the risk of depressive disorders: A population-based study. J. Affect. Disord. 182, 44–49. https://doi.org/10.1016/j.jad.2015.04.027

Chow, C., Epp, J.R., Lieblich, S.E., Barha, C.K., Galea, L.A.M., 2013. Sex differences in neurogenesis and activation of new neurons in response to spatial learning and memory. Psychoneuroendocrinology 38, 1236–1250. https://doi.org/10.1016/j.psyneuen.2012.11.007

Dalla, C., Papachristos, E.B., Whetstone, A.S., Shors, T.J., 2009. Female rats learn trace memories better than male rats and consequently retain a greater proportion of new neurons in their hippocampi. Proc. Natl. Acad. Sci. U. S. A. 106, 2927–32. https://doi.org/10.1073/pnas.0809650106

Falconer, E.M., Galea, L.A.M., 2003. Sex differences in cell proliferation, cell death and defensive behavior following acute predator odor stress in adult rats. Brain Res. 975, 22–36.

Fanselow, M.S., Dong, H.W., 2010. Are the Dorsal and Ventral Hippocampus Functionally Distinct Structures? Neuron 65, 7–19. https://doi.org/10.1016/j.neuron.2009.11.031

Fester, L., Ribeiro-Gouveia, V., Prange-Kiel, J., von Schassen, C., Bottner, M., Jarry, H., Rune, G.M., 2006. Proliferation and apoptosis of hippocampal granule cells require local oestrogen synthesis. J. Neurochem. 97, 1136–1144. https://doi.org/10.1111/j.1471-4159.2006.03809.x

Frokjaer, V.G., Pinborg, A., Holst, K.K., Overgaard, A., Henningsson, S., Heede, M., Larsen, E.C., Jensen, P.S., Agn, M., Nielsen, A.P., Stenbæk, D.S., da Cunha-Bang, S., Lehel, S., Siebner, H.R., Mikkelsen, J.D., Svarer, C., Knudsen, G.M., 2015. Role of Serotonin Transporter Changes in Depressive Responses to Sex-Steroid Hormone Manipulation: A Positron Emission Tomography Study. Biol. Psychiatry 78, 534–543. https://doi.org/10.1016/j.biopsych.2015.04.015

Green, A.D., Galea, L.A.M., 2008. Adult hippocampal cell proliferation is suppressed with estrogen withdrawal after a hormone-simulated pregnancy. https://doi.org/10.1016/j.yhbeh.2008.02.023

Gross, M., Pinhasov, A., 2016. Chronic mild stress in submissive mice: Marked polydipsia and social avoidance without hedonic deficit in the sucrose preference test. Behav. Brain Res. 298, 25–34. https://doi.org/10.1016/j.bbr.2015.10.049

Ingberg, E., Theodorsson, A., Theodorsson, E., Strom, J.O., 2012. Methods for long-term 17β-estradiol administration to mice. Gen. Comp. Endocrinol. 175, 188–93. https://doi.org/10.1016/j.ygcen.2011.11.014

Kokras, N., Pastromas, N., Papasava, D., de Bournonville, C., Cornil, C.A., Dalla, C., 2018. Sex differences in behavioral and neurochemical effects of gonadectomy and aromatase inhibition in rats. Psychoneuroendocrinology 87, 93–107. https://doi.org/10.1016/j.psyneuen.2017.10.007

Kokras, N., Pastromas, N., Porto, T.H., Kafetzopoulos, V., Mavridis, T., Dalla, C., 2014. Acute but not sustained aromatase inhibition displays antidepressant properties. Int. J. Neuropsychopharmacol. 17, 1307–1313. https://doi.org/10.1017/S1461145714000212

Kott, J.M., Mooney-Leber, S.M., Shoubah, F.A., Brummelte, S., 2016. Effectiveness of different corticosterone administration methods to elevate corticosterone serum levels, induce depressive-like behavior, and affect neurogenesis levels in female rats. Neuroscience 312, 201–214. https://doi.org/10.1016/j.neuroscience.2015.11.006

Mahmoud, R., Wainwright, S.R., Chaiton, J.A., Lieblich, S.E., Galea, L.A.M., 2016a. Ovarian hormones, but not fluoxetine, impart resilience within a chronic unpredictable stress model in middle-aged female rats. Neuropharmacology 107, 278–293. https://doi.org/10.1016/j.neuropharm.2016.01.033

Mahmoud, R., Wainwright, S.R., Galea, L.A.M., 2016b. Sex hormones and adult hippocampal neurogenesis: Regulation, implications, and potential mechanisms. Front. Neuroendocrinol. 41, 129–152. https://doi.org/10.1016/J.YFRNE.2016.03.002

Meng, F.T., Ni, R.J., Zhang, Z., Zhao, J., Liu, Y.J., Zhou, J.N., 2011. Inhibition of oestrogen biosynthesis induces mild anxiety in C57BL/6J ovariectomized female mice. Neurosci. Bull. 27, 241–250. https://doi.org/10.1007/s12264-011-1014-8

Molendijk ML, de Kloet ER. Coping with the forced swim stressor: Current state-of-the-art. Behav Brain Res. 2019;364:1–10.

Pawluski, J.L., Van Donkelaar, E., Abrams, Z., Houbart, V., Fillet, M., Steinbusch, H.W.M., Charlier, T.D., 2014. Fluoxetine dose and administration method differentially affect hippocampal plasticity in adult female rats. Neural Plast. 2014. https://doi.org/10.1155/2014/123026

Plümpe, T., Ehninger, D., Steiner, B., Klempin, F., Jessberger, S., Brandt, M., Römer, B., Rodriguez, G.R., Kronenberg, G., Kempermann, G., 2006. Variability of doublecortin-associated dendrite maturation in adult hippocampal neurogenesis is independent of the regulation of precursor cell proliferation. BMC Neurosci. 7. https://doi.org/10.1186/1471-2202-7-77

Saeedi Saravi, S.S., Arefidoust, A., Yaftian, R., Saeedi Saravi, S.S., Dehpour, A.R., 2016. 17α-ethinyl estradiol attenuates depressive-like behavior through GABAA receptor activation/nitrergic pathway blockade in ovariectomized mice. Psychopharmacology (Berl). 233, 1467–85. https://doi.org/10.1007/s00213-016-4242-9

Sherwin, B.B., 1988. Affective changes with estrogen and androgen replacement theraphy in surgically menopausal women. J. Affect. Disord. 14, 177–187. https://doi.org/10.1016/0165-0327(88)90061-4

Snyder, J.S., Choe, J.S., Clifford, M.A., Jeurling, S.I., Hurley, P., Brown, A., Kamhi, J.F., Cameron, H.A., 2009. Adult-Born Hippocampal Neurons Are More Numerous, Faster Maturing, and More Involved in Behavior in Rats than in Mice. J. Neurosci. 29, 14484–14495. https://doi.org/10.1523/JNEUROSCI.1768-09.2009

Strekalova, T., Gorenkova, N., Schunk, E., Dolgov, O., Bartsch, D., 2006. Selective effects of citalopram in a mouse model of stress-induced anhedonia with a control for chronic stress. Behav. Pharmacol. 17, 271–87.

Tuscher JJ, Szinte JS, Starrett JR, Krentzel AA, Fortress AM, Remage-Healey L, Frick KM. Inhibition of local estrogen synthesis in the hippocampus impairs hippocampal memory consolidation in ovariectomized female mice. Horm Behav. 2016;83:60–67.

Tzeng, W.-Y., Chen, L.-H., Cherng, C.G., Tsai, Y.-N., Yu, L., 2014. Sex differences and the modulating effects of gonadal hormones on basal and the stressor-decreased newly proliferative cells and neuroblasts in dentate gyrus. Psychoneuroendocrinology 42, 24–37. https://doi.org/10.1016/j.psyneuen.2014.01.003

Wainwright, S.R., Barha, C.K., Hamson, D.K., Epp, J.R., Chow, C., Lieblich, S.E., Rutishauser, U., Galea, L.A., 2016. Enzymatic Depletion of the Polysialic Acid Moiety Associated with the Neural Cell Adhesion Molecule Inhibits Antidepressant Efficacy. Neuropsychopharmacology 41, 1670–80. https://doi.org/10.1038/npp.2015.337

Wainwright, S.R., Lieblich, S.E., Galea, L.A.M., 2011. Hypogonadism predisposes males to the development of behavioural and neuroplastic depressive phenotypes. Psychoneuroendocrinology 36, 1327–1341. https://doi.org/10.1016/j.psyneuen.2011.03.004

Yagi, S., Chow, C., Lieblich, S.E., Galea, L.A.M.M., 2016. Sex and strategy use matters for pattern separation, adult neurogenesis, and immediate early gene expression in the hippocampus. Hippocampus 26, 87–101. https://doi.org/10.1002/hipo.22493

Zarrouf, F.A., Artz, S., Griffith, J., Sirbu, C., Kommor, M., 2009. Testosterone and depression: systematic review and meta-analysis. J. Psychiatr. Pract. 15, 289–305. https://doi.org/10.1097/01.pra.0000358315.88931.fc

